# Analog intrinsic recoding measures RNA dynamics without chemical conversion

**DOI:** 10.64898/2026.07.27.741041

**Authors:** Lisa N. Hansen, Mary T. Couvillion, Erik McShane, Nadezhda Azbukina, Barbara Treutlein, L. Stirling Churchman

## Abstract

Steady-state RNA abundance measurements mask the synthesis and decay rates that shape gene expression. Analog intrinsic recoding sequencing (AIR-seq) repurposes the base-pairing properties of N4-hydroxycytidine (NHC) to mark newly synthesized RNA with C-to-T and T-to-C mismatches in standard RNA-seq libraries, eliminating chemical conversion and enrichment common to RNA metabolic labeling methods. NHC-AIR-seq resolves bulk and single-cell RNA dynamics, improves RNA velocity inference, reveals hidden regulation, and brings RNA kinetics into routine transcriptomic workflows.

## Main text

Steady-state RNA abundance reflects the combined effects of synthesis, processing, export, and decay, meaning distinct kinetic regimes can produce indistinguishable steady-state levels. Conventional metabolic-labeling RNA-seq methods, including nucleotide-recoding RNA-seq (NR-seq; **Fig. 1a**), can resolve these processes, but require chemical-recoding steps that can complicate workflows, damage RNA, and are poorly compatible with standard droplet-based single-cell genomics^1–3^. Here we introduce analog intrinsic recoding sequencing (AIR-seq), which uses β-D-N4-hydroxycytidine (NHC) to mark newly synthesized RNA without the post-labeling chemistry used by nucleotide recoding sequencing (**Fig. 1a**). NHC, the active compound of the COVID-19 antiviral molnupiravir, can be incorporated into cellular RNA by RNA polymerase II and undergoes ambiguous base pairing^4^. We therefore reasoned that reverse transcription of NHC-containing RNA would generate C-to-T and T-to-C mismatches in otherwise standard RNA-seq libraries, eliminating both enrichment and conversion chemical steps required by existing approaches.

**Figure 1:**
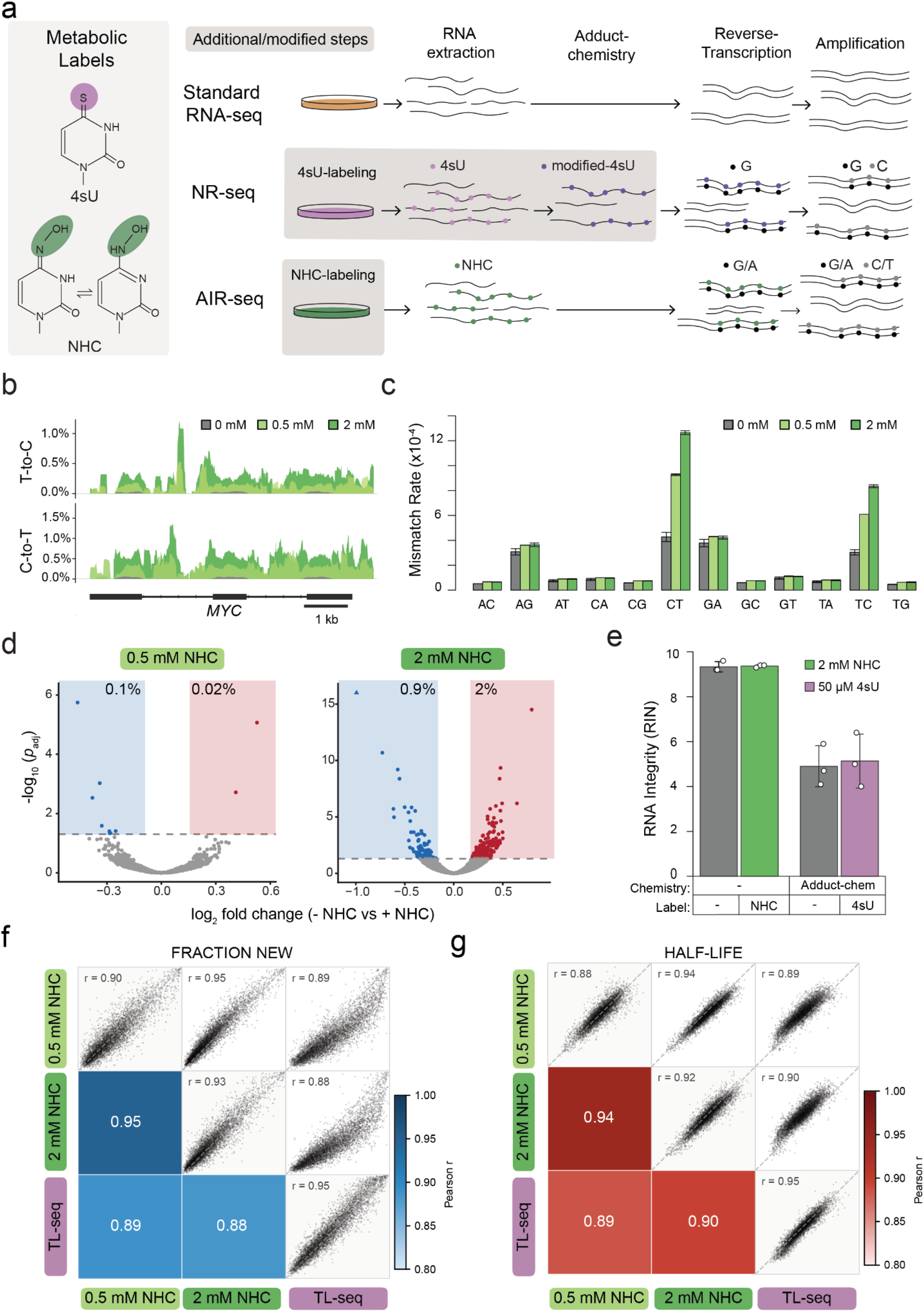
NHC-AIR-seq measures RNA stability. All experiments in HeLa cells, 2 h treatment with 0, 0.5, or 2 mM NHC (4sU for TimeLapse-seq (TL-seq) comparisons), unless noted. **a)** Structures of 4sU and the NHC tautomer (left panel); workflows for standard RNA-seq, NR-seq, and AIR-seq (right panel). **b)** T-to-C and C-to-T mismatch rate along the *MYC* locus. Rate = mismatched reads/ total reads per position, smoothed (150 bp rolling mean, gaps > 30 bp left unsmoothed; positions with >3x the sample median masked as putative SNPs). Gene model below shows exon/intron structure, scale bar 1 kb. **c)** Mismatch rates for all 12 possible single-nucleotide substitution types in HeLa exonic reads (mean ± range, n = 2 replicates). **d)** Differential expression by NHC dose vs unlabeled control (left, 0.5 mM; right, 2 mM). DESeq2 (Wald test; log2 fold change shrinkage ‘normal’ prior). Red, up-regulated; blue, down-regulated; grey, not significant (padj<0.05, Benjamini-Hochberg FDR correction). Triangle indicates PPP1R10, plotted at the y-axis maximum for visibility; percentages indicate fraction of transcriptome affected. **e)** RNA integrity (RIN) of NHC- and TL-seq ± metabolic label. RNA used for library prep, relative to matched untreated controls (4sU chemically converted; NHC not). Bars, mean ± s.d.; points, individual reps (n = 3). **f)** Pairwise agreement in fraction-new among NHC-AIR-seq vs. TL-seq. Lower triangle, Pearson r (replicates pooled); diagonal, replicate A vs B (r shown); n=5,804 genes (shared read depth > 100 in all samples). **g)** As in **(f)**, for RNA half-life, n=5,800 genes, to the same read depth as in fraction new and half-life within a plausible range (0.1-100 hr).

To establish NHC-AIR-seq, we treated HeLa and HEK293T cells with two NHC concentrations and performed standard paired-end RNA-seq (HeLa, **Fig. 1**; HEK293T **Supplementary Fig. 1b-e**). Both C-to-T and T-to-C mismatch rates rose with NHC concentration and varied by polymerase, with minimal transcriptome perturbation (**Fig. 1b-d, Supplementary Fig. 1a-c**). NHC also preserved RNA integrity (RIN 9.2-9.6), in contrast to periodate conversion used by the NR-seq method TimeLapse-seq^5^ (RIN 4.0-6.4; **Fig. 1e**). Using a customized version of the EZbakR^6^ pipeline that maximizes signal-to-noise and uses both C-to-T and T-to-C conversions, we estimated the fraction of new RNA (fraction-new) transcriptome-wide. Estimates were reproducible across replicates and NHC concentrations and concordant with TimeLapse-seq^5^ for fraction-new and RNA half-life estimates (Pearson’s r>0.84; **Fig. 1f-g and Supplementary Fig. 1d-e**). To define practical sequencing requirements, we downsampled across paired-end, single-end and 3′ libraries and found that these results were reproducible across a wide range of sequencing depths and library preparation types (**Supplementary Fig. 1f-g**).

Single-cell RNA dynamics measurements are especially constrained by sparse RNA recovery, which leaves little tolerance for the losses introduced by enrichment or chemical conversion. To establish the performance of NHC-AIR-seq in single cells, we labeled HeLa cells for 6 h and processed them using the standard 5′ v2 10x Genomics workflow. Using both T-to-C and C-to-T mutations, we estimated fraction-new for 2,416 genes across 1,610 cells with a one-parameter binomial mixture model calibrated using the NHC incorporation rate inferred by pseudobulk EZbakR. The resulting posterior read probabilities were used to derive new- and old-RNA count matrices alongside the observed total-RNA counts. Median single-cell estimates closely matched pseudobulk values (Pearson’s r = 0.98; **Fig. 2a**) and remained concordant after reads were truncated to standard read lengths in single-cell applications (Pearson’s r = 0.93 and 0.79 for 150 or 90 nt respectively; **Supplementary Fig. 2a**). NHC-AIR-seq gene detection scales with read depth (**Supplementary Fig. 2b**). After correcting for sampling noise, cell-cycle genes showed greater cell-to-cell fraction-new variability than housekeeping genes, consistent with dynamic RNA synthesis and decay across the cell cycle (**Supplementary Fig. 2c**).

**Figure 2:**
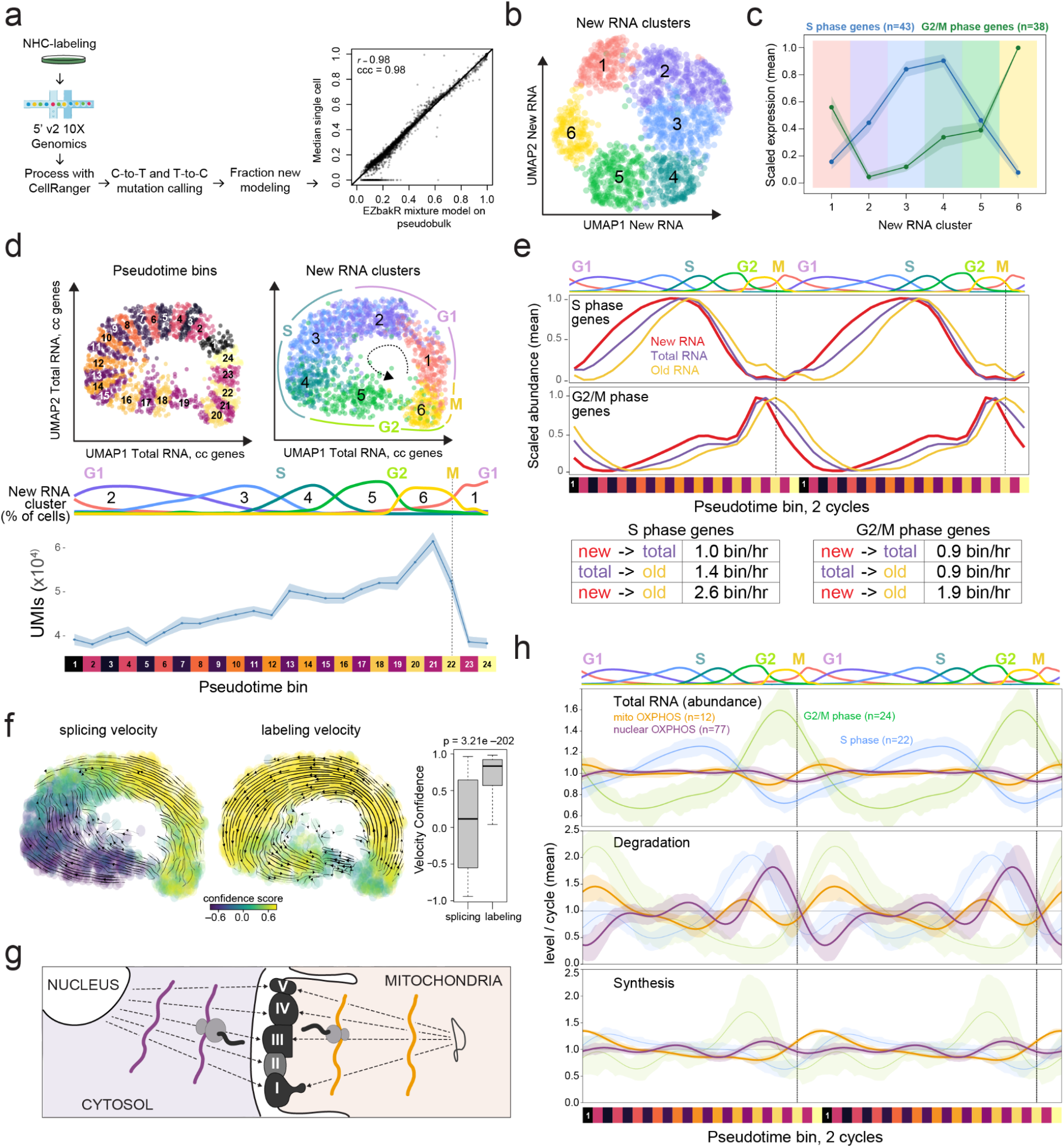
Single-cell NHC-AIR-seq resolves RNA dynamics across the cell cycle. **a)** Single-cell NHC-AIR-seq schematic workflow and correlation between median fraction-new values and pseudobulk EZbakR fraction-new values for 2416 genes. **b)** UMAP for the top 2,000 variable genes in new-RNA. **c)** Mean scaled expression of S-phase and G2M-phase genes for each new-RNA cluster +/-95% confidence interval. **d)** UMAP for 196 cell-cycle genes in total-RNA split into 24 equal-cell bins along pseudotime, starting at total-RNA G1 phase (see Supplementary Fig 2f). New-RNA variable-gene clusters are labeled along with inferred cell cycle phase. Arrow shows direction of cell cycle progression. Bottom: mean UMI counts/cell in each pseudotime bin +/-standard error. Dotted line indicates cell division. **e)** Smoothed mean expression across pseudotime bins. Temporal offsets determined by cross-correlation analysis, with one bin estimated as one hour. **f)** RNA velocity estimated from splicing dynamics and metabolic labeling dynamics using Dynamo, on a matched gene panel consisting of 82 cell cycle genes. Streamlines show the consensus velocity field colored by velocity confidence score. Wilcoxon p is shown. **g)** Schematic of the five dual-origin OXPHOS complexes in the mitochondrial inner membrane. **h)** Modified FourierCycle estimates of total RNA abundance, degradation, and synthesis (top to bottom) across pseudotime. Each gene was normalized to its mean across the cell cycle and then averaged within gene sets; shading indicates interquartile ranges. Two cycles are displayed; the dotted line marks cell division (bin 22).

We first tested whether fraction-new variability reflected cell state. Cells formed a ring-like manifold in the uniform manifold approximation and projection (UMAP) embeddings derived from new-RNA counts. Gene Ontology (GO) and the relative abundances of canonical S-phase and G2/M programs clearly ordered clusters through G1, S, late S/G2 and G2/M phases (**Fig. 2b,c and Supplementary Fig. 2d,e**). Total RNA recovered a related structure but less clearly separated G1 cells (**Supplementary Fig. 2e**).

To resolve the temporal organization of these states, we constructed a cell-cycle UMAP from total-RNA expression of cell-cycle genes and converted position along the trajectory to 24-bin pseudotime anchored at the onset of G1 in total RNA. Mean unique molecular identifier (UMI) counts approximately doubled before dropping at division, as observed in other proliferating single-cell datasets^7^. The drop coincided with the transition from new-RNA cluster 6 to cluster 1, orienting the inferred cell-cycle phases along the trajectory (**Fig. 2d**). Across pseudotime, S-phase and G2/M programs appeared first in new RNA, followed by total and then old RNA (**Supplementary Fig. 2f**). Circular cross-correlation quantified a ∼1-h lead for new RNA and a 0.9–1.4-h lag for old RNA, separating new and old profiles by approximately 2 hours (**Fig. 2e**). As an independent test, Dynamo^8^ velocities derived from new and total RNA showed greater local directional consistency and stronger alignment with pseudotime than velocities inferred from conventional splicing ratios (**Fig. 2f**). These advantages persisted after reads were trimmed to single-end lengths of 150 or 90 nt (**Supplementary Fig. 2g**). Thus, a single NHC-AIR-seq snapshot resolves cell-cycle RNA dynamics at hourly resolution and improves trajectory inference without requiring a population time course.

We next investigated whether single-cell rate modeling could reveal regulation masked by stable RNA abundance. We focused on oxidative phosphorylation (OXPHOS), whose complexes combine mitochondrial- and nuclear-encoded subunits that must be coordinately expressed despite distinct bulk kinetics^5^ (**Fig. 2g**). We adapted FourierCycle^8^ to NHC-AIR-seq-derived new and total RNA. Although nuclear and mitochondrial OXPHOS mRNAs showed little oscillation in abundance (**Fig. 2h**, top), inferred synthesis and degradation rates revealed divergent cell-cycle programs. Nuclear transcripts showed a degradation pulse before division, followed by low degradation in early G1, whereas mitochondrial transcripts showed an early-G1 pulse of synthesis and degradation (**Fig. 2h**, middle and bottom). These offset programs could renew nuclear OXPHOS mRNAs as mitochondrial OXPHOS synthesis rises in early G1, coordinating expression across the two genomes without large changes in abundance. Thus, NHC-AIR-seq reveals kinetic regulation that is invisible in steady-state RNA.

AIR-seq blows past the chemical-conversion and enrichment bottlenecks of current metabolic-labeling approaches, enabling RNA-dynamics measurements in standard bulk and single-cell workflows. The clinical use of its prodrug molnupiravir raises the possibility of *in vivo* RNA-dynamics measurements, while other intrinsically recoding analogs may offer alternative implementations of AIR-seq. Therefore, AIR-seq establishes a simple and broadly compatible approach for quantitative measurements of RNA dynamics in standard bulk and single-cell sequencing workflows.

## Methods

### Materials

All commercially available materials were purchased from the indicated suppliers and used without further purification. Beta-d-N4-Hydroxycytidine (NHC) was purchased from Cell Signaling Technology (Danvers, Massachusetts). Gibco Dulbecco’s Modified Eagle Medium (DMEM), Gibco Fetal Bovine Serum, Premium (FBS), Gibco phosphate buffered saline (PBS), Invitrogen Sodium Acetate (C_2_H_3_NaO_2_), Invitrogen Ambion GlycoBlue Coprecipitant, Trizol LS reagent, dithiothreitol (DTT), and Penicillin Streptomycin (10,000 U/mL) were purchased from ThermoFischer Scientific (Waltham, Massachusetts). DNase I was purchased from New England Biolabs (Ipswitch, Massachusetts). Takara SMART-Seq Total RNA Pico Input with UMIs (ZapR Mammalian). AMPure XP Beads and RNAClean XP RNA and cDNA Cleanup Reagent were purchased from Beckman Coulter (Brea, California). DNA/RNA Shield was purchased from Zymo (Irvine, California). HeLa cells and Universal Mycoplasma Detection Kit were obtained from ATCC (Manassas, Virginia). 4-thiouridine, sodium periodate, and 2,2,2-trifluoroethylamine were obtained from Sigma-Aldrich (St. Louis, Missouri).

### Cell growth

HeLa and HEK293T cells were maintained at 37°C and 5% CO_2_ in DMEM with 10% FBS and 1% P/S. Cells were tested monthly for Mycoplasma with the Universal Mycoplasma Detection Kit (ATCC).

### NHC treatment

80% confluent cells in 12-well plates were labeled with NHC. For bulk RNA-seq, NHC was prepared as 125 mM and 500 mM stock solutions in DMSO and added to final concentrations of 0.5 or 2 mM, respectively. For 3′ bulk RNA-seq and single-cell RNA-seq, NHC was prepared as a 50 mM stock solution in water and added to a final concentration of 2 mM. Corresponding DMSO or water vehicle controls were included for each experiment. At the beginning of each labeling period, cells were removed from the incubator, NHC was added directly to existing cell media, and cells were returned to the incubator for the remainder of the pulse. At the end of the pulse, the cell media was removed and discarded, and cells were immediately put on ice and lysed in 500 µL RIPA buffer to collect total RNA.

### RNA extraction and DNase treatment

RNA extraction was performed using Trizol LS according to the manufacturer’s protocol. RNA was quantified by Nanodrop. RNA was treated with DNase I for 30 min at 37 °C and cleaned with 2× volume of RNA clean beads and 2 washes with 80% ethanol.

### Library preparation

Library preparation was performed using the SMART-Seq Total RNA Pico Input with UMIs (ZapR Mammalian) with 10 ng of RNA. Samples were sequenced on a NovaSeq using paired-end 2×150 reads to a median depth of 56 million reads (range 41-110 million) per sample by the Biopolymers Facility at Harvard Medical School.

### RNA integrity

90% confluent HeLa cells were treated with 2 mM NHC or 50 µM 4sU for 2 hours. RNA was extracted as above with the addition of 0.1 mM DTT for 4sU samples, and all 4sU samples were handled in the dark. Samples were DNase-treated, and then 4sU-treated samples and controls underwent TimeLapse chemistry as detailed in Schofield et al^9^. Samples were then submitted to the Biopolymers Facility at HMS for TapeStation.

### Submitting NHC samples for 3′end RNA-seq

Cells were treated with NHC, as described above, for 2 hours with 2 mM NHC. Following treatment, cells were removed from the incubator, media was removed, and cells were washed 1x with PBS and trypsinized. Cells were counted, and 2.5 × 10^5^ cells were suspended in 50 µL of Zymo DNA/RNA shield. RNA-seq libraries were prepared and sequenced by Plasmidsaurus (Pleasanton, CA, USA) using a stranded 3′-end counting workflow. Poly(A) RNA was captured by oligo(dT)-primed reverse transcription with incorporation of sample barcodes and unique molecular identifiers (UMIs), followed by second-strand synthesis, tagmentation, library amplification with unique dual indices, and Illumina sequencing to a target depth of up to 10 million deduplicated reads per sample with read length ∼90 bp aligned within the last ∼400nt of the transcript.

### fastq2EZbakR and EZbakR

Raw reads in fastq format were processed into a counts binomial file (number of T-to-C/C-to-T mismatches and number of T/C per read) using the fastq2EZbakR pipeline ^2^ with several modifications necessary to improve signal-to-noise in NHC-treated vs untreated samples. First, reads are adaptor-trimmed and hard-clipped before entering the pipeline. This step is needed to prevent alignment-artifact mismatches at ends caused by partial matching within the random hexamer or template-switch regions. Next, an option is added to remove reads that map to mitochondrial RNA polymerase and common RNA polymerase III transcripts. These enzymes appear to more readily incorporate NHC, resulting in higher mutation rates, which provide spurious information to the mixture model for PolII transcripts (**Supplementary Fig. 1a**). Finally, SNP-calling was changed from true SNP-calling using genotype likelihoods to using simple counting in untreated samples, with the reasoning that any position that is commonly mismatched in a control sample processed in parallel is not trustworthy for calling NHC-induced mismatches. The modified pipeline also retains cell barcode and other single-cell-relevant information when available.

In addition to the pipeline modifications, parameters were also optimized to minimize background mutations. Notably, alignment with STAR to the human reference genome (GRCh38/hg38) uses stricter thresholds than default, and the minimum base quality for calling a mismatch is set at one less than the maximum quality, though on-machine quality binning for our bulk runs (Novaseq X Plus) limits the utility of this setting (e.g. any value ≥ Q30 receives the score Q40).

Counts binomial files were processed in R using EZbakR (https://github.com/isaacvock/EZbakR) to estimate new-to-total ratio (NTR/fraction-new) and kdeg for each transcript. In order to capture only two distinct populations (high mutation content [labeled] and low mutation content [unlabeled]), T-to-C mismatches and C-to-T mismatches were summed per read, as were the number of T and C. Intronic reads were presumed to originate from unspliced transcripts and were not considered.

For comparisons between NHC-AIR-seq and TimeLapse-seq (fraction-new and half-life agreement analyses), only genes with >100 uniquely mapped reads in all compared samples were used. For half-life comparisons specifically, genes were further restricted to those with half-life estimates within a well-resolved range (0.1-100 h) in all compared conditions, excluding values poorly identifiable within the 2 h labeling window.

### TimeLapse-seq re-analysis

Publicly available HeLa TimeLapse-seq data ^5^ were reprocessed from raw sequencing reads using the same fastq2EZbakR/EZbakR pipeline and parameters described above. HEK293T TL-seq data^10^ were not reprocessed here; half-life estimates were obtained from Vock et al^2^, where the original data was reprocessed using the fastq2EZbakR/EZbakR pipeline, with replicates pooled (**Supplementary Fig. 1e**).

### Differential gene expression analysis

Differential gene expression between NHC-labeled and unlabeled HeLa or HEK293T cells (0, 0.5, and 2 mM NHC, 2 h labeling; n=2 biological replicates per dose) was assessed using DESeq2 (v1.50.2) in R. Gene-level counts were derived from fastq2EZbakR output, with mitochondrial and PolIII genes excluded either during fastq2EZbakR with a bam of mito/PolIII (HeLa cells) or by filtering the fastq2EZbakR output with a gtf based mito/PolIII list (HEK293T). Genes with fewer than 10 reads in fewer than 2 samples were excluded prior to testing. Counts from all three doses were fit jointly under a single design (∼dose), and each labeled dose was compared to the unlabeled (0 mM) control via a Wald test, with log2 fold changes shrunk using a normal estimator^11^. Genes with a Benjamini-Hochberg-adjusted p < 0.05 were considered significantly differentially expressed.

### Downsampling

To assess the effect of sequencing depth on NHC-AIR-seq estimates, we performed *in silico* downsampling on the compressed read count files (cB files) generated by fastq2EZbakR. For single-end analysis, read 2 from the paired-end experiment was used as input to fastq2EZbakR in single-end mode with strandedness set to forward. In this reverse-stranded library preparation, read 2 represents the sense strand of the original RNA; correct strand orientation was confirmed by the presence of T>C and C>T mutations consistent with NHC incorporation in labeled samples. For each library type (paired-end 2×150bp, single-end 150bp, and 3′-end 90bp), read counts per gene were resampled to 5%, 10%, 25%, 50%, 75%, and 90% of the original depth, as well as full depth as a reference, using multinomial sampling, preserving the relative gene expression distribution. Fraction-new was re-estimated at each depth using EZbakR with identical parameters to the full-depth analysis. Across seeds, Pearson correlation with the full-depth reference was averaged via Fisher z-transformation (weighted by degrees of freedom), with 95% confidence intervals computed in z-space and back-transformed; gene retention (% of full-depth genes with a valid estimate at each depth) was averaged as mean ± s.d. Correlation estimates from cells representing fewer than 20 genes in any replicate were flagged as unreliable; no cell in the final dataset fell below this threshold. Genes were stratified into expression quartiles based on total exonic read counts in the full-depth paired-end dataset; these boundaries were applied to single-end data, for which gene ranking is preserved by construction (single-end reads are one mate of the same paired-end read pairs), and to 3′-end data, a distinct library chemistry from a separate sequencing run for which expression ranking may not be fully concordant with paired-end coverage. The paired-end and single-end datasets comprised a mean of 42.9 million exonic reads per sample; the 3′-end dataset, 14.1 million.

### Single-cell NHC-AIR-seq

HeLa cells were treated with 2 mM NHC for 6 hours as in bulk and were harvested by trypsinization, quenched with DMEM/10% FBS, and washed twice in PBS + 0.04% BSA. Cell suspensions were filtered through 40 µm strainers, counted, and adjusted to 700–1,200 cells/µL. The library was generated using the 10x Genomics Chromium Next GEM 5′ v2 kit (protocol CG000331 Rev F) targeting 2,000 cells, followed by gene expression (GEX) library construction with paired-end, dual-indexed sequencing. The library was sequenced on an Element Biosciences AVITI instrument (Low Output, 250M reads, 2×150 kit), yielding 76,401 mean reads per cell. For analysis of 150 nt and 90 nt single-end reads, the Read 1 sequence was trimmed *in silico* to 26 nt to include only the cell barcode and UMI, and the Read 2 sequence was trimmed to 150 or 90 nt respectively.

### Analysis of single-cell NHC-AIR-seq

Single cell NHC-AIR-seq data was analyzed by integrating the existing tools CellRanger and fastq2EZbakR (bam2EZbakR variant), as well as custom scripts for estimating fraction new per gene in single cells. In detail, fastq files were processed with 10X Genomics Cell Ranger v9.0.1^12^ to align, filter, and count barcodes (cells) and UMIs (molecules). Cell Ranger uses STAR to align reads with parameters that could not be adjusted to our optimized stringent settings. However, two differences in the AVITI instrument used here compared to Novaseq used for bulk sequencing help to decrease background. First AVITI Low Output results in higher overall quality scores (>90% Q30 vs ≥85% Q30). Second, quality scores are not binned, meaning bases with even moderately reduced qualities (Q30 - Q38) can be removed from consideration. The resulting alignment file (.bam) is then formatted for the bam2EZbakR pipeline with samtools by adding the ‘string for mismatching positions’ (MD) tag. Our modified fastq2EZbakR carries the cell barcode information as well as Cell Ranger’s exon/intron/intergenic assignment through to the final counts binomial file (cB.csv). Note that adaptors are still present after Cell Ranger alignment resulting in soft-clipped reads extending beyond transcript ends and causing EZbakR to fail to correctly designate exonic reads. Thus we use Cell Ranger’s exon assignment instead.

To estimate per-cell fraction new values for genes passing the expression threshold (>= 10 reads in at least 1% of cells), we used a calibrated method-of-moments estimator that borrows the inferred mutation rate (p_*new*_: conversion probability of newly synthesized reads) from the EZBakR Maximum Likelihood Estimation (MLE) mixture model fit performed on pseudobulk data. For each read, *k* denotes the number of observed conversion events and *n* the number of convertible nucleotide positions. To improve robustness in sparse single-cell data, reads were binarized according to whether at least one conversion event was detected (*k* > 0), rather than modeling the full conversion count distribution. The expected probability of detecting at least one conversion in newly synthesized reads was modeled as 1−(1−p_*new*_)^*n*^. Cell-level fraction new values were estimated by matching the observed fraction of reads containing at least one conversion to the expected mixture of new and pre-existing reads. For each gene, the background conversion rate in pre-existing RNA was estimated from control reads *across all cells* as the total number of observed conversions divided by the total number of convertible sites. We then computed posterior probabilities that each read originated from newly synthesized RNA using Bayes’ rule and averaged these probabilities to obtain the final fraction-new estimate.

Standard deviations in fraction new across cells were noise-corrected by modeling each per-cell estimate as a binomial proportion and subtracting the expected sampling variance. For a given gene, the fraction new in cell i, fni, is estimated from N_i_ reads (the per-cell read count for that gene) and is therefore subject to binomial sampling noise of variance fn_i_(1 − fn_i_)/N_i_, which is larger in cells with fewer reads. The variance in fraction new observed across cells thus reflects the sum of true biological cell-to-cell variation and this sampling noise. Because variances are additive, we estimated the biological component by a method-of-moments correction, subtracting the mean per-cell sampling variance from the observed across-cell variance.

### Reductions and clustering

For total-RNA UMAP and new-RNA UMAP, the top 2,000 most variable genes in each assay were identified using the Seurat v5.4.0^13^ default variance-stabilizing selection, scaled, and used as input to PCA. The first 12 or 20 principal components for new-RNA and total-RNA respectively were used to construct a shared nearest-neighbor graph (Leiden algorithm, resolution set for 6 clusters) for clustering. For the Cell Cycle UMAP, inputs were restricted to a curated set of 196 cell-cycle genes. Expression values for these genes were scaled and subjected to principal component analysis (PCA). The first 20 principal components were used as input for UMAP visualization.

Cell-cycle phase was scored using a manually curated, HeLa-oriented gene set comprising 43 genes strongly associated with replication-associated S-phase and 39 mitotic G2/M-phase genes based off (Whitfield ML et al. 2002. Mol Biol Cell^14^). Seurat CellCycleScoring was used to calculate S and G2/M scores and assign an initial phase. A module score was also calculated for replication-dependent histone genes expressed in this dataset. Cells initially classified as G1 were reassigned to S or G2/M when the histone score was positive, according to whether the S or G2/M score was greater; all other assignments were retained.

For Gene Ontology semantic-reduction heatmaps, Biological Process enrichment was performed separately for clusters defined from total-RNA and new-RNA expression profiles using the enrichGO function of the clusterProfiler R package (v4.14.6)^15,16^. GO term annotations were drawn from the human genome-wide annotation database org.Hs.eg.db (Bioconductor, v3.20.0), with marker genes matched by official gene symbol. For each cluster, significantly upregulated marker genes were selected using a Benjamini–Hochberg-adjusted (P)-value threshold of 0.01, an average absolute log2 fold-change of at least 0.15, and expression in at least 10% of cells within the cluster. Terms with adjusted P<0.05 were retained, and the 25 most significant terms per cluster, ranked by -log_10_ adjusted P-value, were used for visualization. Redundant GO terms were grouped using semantic similarity analysis implemented in rrvgo (v1.18.0)^17^. Pairwise term similarities were computed with the Rel method (GOSemSim, v2.32.0)^18^, and terms were collapsed into representative parent themes using a similarity threshold of 0.7. Each GO term was assigned to the representative parent theme identified in the cluster in which that term had its strongest enrichment. Parent themes and their constituent terms were ordered according to the cluster in which they reached their maximum enrichment, followed by decreasing maximum -log_10_ adjusted P-value.

### Pseudotime

Cell cycle pseudotime was computed from a UMAP embedding of total-RNA. A center point was estimated by minimizing the median absolute deviation of cell-to-center radii, and each cell was assigned an initial angle using the two-argument arctangent (atan2). To correct for non-circularity of the UMAP loop, a smooth closed curve was fit to the trajectory using smoothing splines (smooth.spline, R stats package) on 200 angular bins, and cells were re-parameterized by cumulative arc length along this curve, each cell being assigned the arc-length position of its nearest point on the curve. The resulting pseudotime was rotated so that it starts at the border of G2/M and G1 clusters as determined by Seurat CellCycleScoring on total-RNA (see Supp Fig 2f, middle panel) and oriented so that pseudotime increases in the order G1 → S → G2M. For visualization of metadata and gene expression trends, cells were ranked by arc-length pseudotime and divided into 24 bins containing approximately equal numbers of cells. Within each bin, values were averaged across cells to generate pseudotime profiles.

### Cross-correlation offset

For each assay (new-RNA, total-RNA, and old-RNA), normalized expression was averaged across cells within each bin for predefined S- and G2/M-phase gene sets. Gene-set profiles were standardized within each assay and smoothed using a circular three-point weighted moving average (weights 0.25, 0.50, and 0.25), with circular padding to preserve continuity across the cell-cycle boundary. Smoothed profiles were plotted together to compare the temporal progression of gene expression across assays. To quantify global temporal shifts between assays, circular cross-correlation was computed on the smoothed expression profiles after z-score normalization. Profiles were upsampled by linear interpolation to achieve sub-bin resolution, and the lag maximizing the normalized cross-correlation coefficient was reported as the relative delay between assays.

### RNA velocity with Dynamo

RNA velocity was estimated from both metabolic labeling and RNA splicing using Dynamo on a matched panel of 82 cell-cycle genes present in both sets and a shared set of cells. For splicing-based velocity, spliced and unspliced transcript counts were generated from Cell Ranger BAM files using velocyto.py run10x with GENCODE gene annotations and RepeatMasker annotations to mask repetitive elements. Metabolic-labeling velocity was estimated from total- and new-RNA count matrices generated from the NHC-AIR-seq labeling analysis. Dynamo was run using the deterministic two-step model with one-shot metabolic labeling (NTR_vel = TRUE) to estimate labeling-based velocities, whereas the conventional deterministic splicing model was used for splicing-based velocities.

Velocity confidence was quantified using two complementary measures: neighbor consistency, defined as the mean cosine similarity between a cell’s velocity vector and those of its nearest neighbors, and arc-tangent alignment, defined as the cosine similarity between the velocity vector and the tangent of the arc-length cell-cycle pseudotime trajectory. The product of these measures was used as a combined confidence score, such that high-confidence velocities were both locally consistent with neighboring cells and aligned with the inferred direction of cell-cycle progression.

### Fourier modeling of cell-cycle–resolved synthesis and degradation

We recovered continuous, cell-cycle–phase–resolved transcription and degradation rates for each gene by adapting the FourierCycle framework^8^ (https://github.com/MolinaLab-IE/FourierCycle), which was developed to infer periodic kinetic rates from single-cell unspliced/spliced RNA. Our implementation retains FourierCycle’s core representation — periodic kinetic rates expanded as truncated Fourier series on the cell-cycle phase axis θ ∈ [0, 1), with rate recovery performed analytically in the Fourier domain — but replaces its splicing-based observation model with one appropriate to metabolic labeling data.

Biophysical model: We model the cell cycle as a phase variable θ ∈ [0, 1), a circular (periodic) coordinate, advancing at rate *d*θ/*dt* = 1/*T*_*cc*_, with cycle period *T*_*cc*_ = 24 h (assumed). For each gene, total mRNA abundance *R*(θ) obeys a single mass-balance ODE:

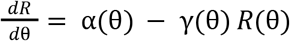

Where α(θ) = *T* _*cc*_ *K* _*syn*_ (θ) and γ(θ) = *T* _*cc*_ *K* _*deg*_ (θ) are the per-cycle synthesis and degradation rates, both periodic with period 1. Whereas FourierCycle requires two coupled state equations — an unspliced species *u* and a spliced species *s*bridged by a fitted scalar splicing rate β that serves as an internal clock — metabolic labeling supplies this clock directly through the known labeling duration *t* _*L*_, collapsing the system to the single equation above.

Observation model for labeling data: A cell sampled at phase θ was labeled over the window [θ − Δ _*L*_, θ] (mod 1), where Δ _*L*_ = *t* _*L*_ /*T* _*cc*_ = 6/24 = 0. 25. Defining the survival factor 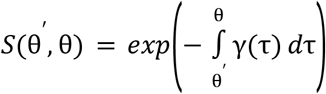, the observables are

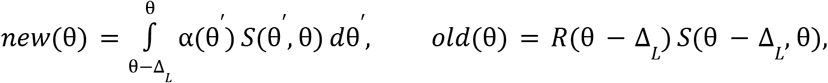

With *total*(θ) = *new*(θ) + *old*(θ) = *R*(θ) by mass conservation across the window. The measured fraction-new is therefore

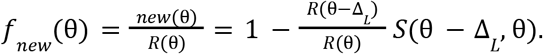

Rearranging yields the window-averaged degradation rate directly from the data:

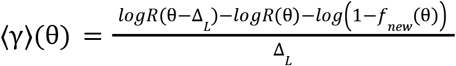

where 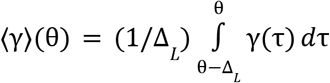 is the boxcar-averaged γ over the labeling window. Critically, because Δ _*L*_ = 0. 25 is not small relative to the cycle, and because *R* oscillates, the *R*(θ − Δ_*L*_)/*R*(θ) prefactor is retained; the naïve estimator − *log*(1 − *f* _*new*_)/Δ _*L*_ conflates degradation modulation with abundance modulation.

Per-gene fitting: Modeling was restricted to the 2,413 genes with new, old, and EZBakR posterior fraction-new estimates, across 1,608 labeled cells (a median of 1,494 cells per gene contributed after per-gene read filtering). Total mRNA counts were library-size normalized (counts × 10^4^ / per-cell total) prior to fitting, removing per-cell depth variation. Each cell carried a cell-cycle pseudotime (cc_pseudotime_arc ∈ [0, 1)), which was used directly as the phase θ. For each gene:

1. **Smoothing**. Cells were sorted by θ and partitioned into 24 equal-cell bins; per-bin means of the fraction-new and total counts were computed. The Fourier basis was parameterized in the rank coordinate *u* = *rank*(θ)/*N*, so bin centers are exactly uniform, and curves were fit directly at the resolved resolution.
2. **Fourier fit**. *R*(θ) and the boxcar estimate ⟨γ⟩(θ) were each fit by least squares to complex Fourier series truncated at *N*_*t*_ = 3 harmonics. Derivatives *dR*/*d*θ were taken analytically (multiplication by 2*πin* in the Fourier domain).
3. **Deconvolution of the labeling window**. The finite window imposes a boxcar (moving-average) blur on ⟨γ⟩ with transfer function *H* _*n*_ = *exp*(− *iπn* Δ _*L*_) *sinc*(*n* Δ _*L*_). Instantaneous γ(θ) was recovered by Wiener-regularized deconvolution,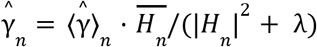, with *λ* = 0. 1; harmonics at the sinc null (|*H* _*n*_ | < 0. 05) were dropped.
4. **Synthesis rate**. α(θ) was computed from the model equation as 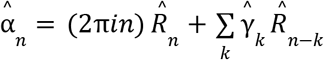 (convolution in the Fourier domain, following FourierCycle’s rate-recovery construction). An *n*^2^-weighted low-pass (ρ = 0. 15) was applied to the copy of *R* feeding α only, to suppress amplification of high-harmonic abundance noise by the derivative.
5. **Non-negativity**. Where α(θ) or γ(θ) went negative on the grid, the nearest non-negative solution was obtained by constrained refit (BFGS), initialized from the closed-form estimate.

Per gene we reported the Fourier coefficients of α, γ, γ _*box*_, and *R*; mean synthesis and degradation rates (h^−1^); oscillation amplitudes, dominant harmonics, and phases; and mRNA half-lives (from mean γ).

Resolution limit: Because the boxcar transfer function first vanishes at *n* = 1/Δ_*L*_ = 4, harmonics |*n*| ≥ 4 are unresolvable; fitting them renders deconvolution noise as spurious high-frequency oscillations, most severely in α (whose derivative term amplifies harmonic *n* by 2*πn*). We therefore fit at *N* _*t*_ = 3, the resolvable band. This is analogous to — but distinct from —FourierCycle’s resolution limit, which is set by the splicing rate β rather than by a labeling window.

Interpretation of kinetic rates: Library normalization removes much of the global increase in RNA content through the cell cycle and its reduction at division, and fraction-new is invariant to this scaling. However, synthesis associated with expansion of the cellular RNA pool can reduce the fraction of pre-existing RNA even without biochemical degradation. Broad phase-dependent changes in γ(θ) is interpreted as relative changes in RNA degradation, while recognizing that library normalization and finite-labeling-window deconvolution limit inference of absolute rate magnitudes and exact peak positions. Biological conclusions consequently focus on broad differences in kinetic timing and profile shape across gene sets.

## Acknowledgements

We would like to thank I. Vock for advice on fastq2EZbakR and EZbakR, E. Duffy, I. Patop, and J. Ly for careful reading of the manuscript. Funding was provided by R01HG007173 to L.S.C.

---

**Supplementary Figure 1:**
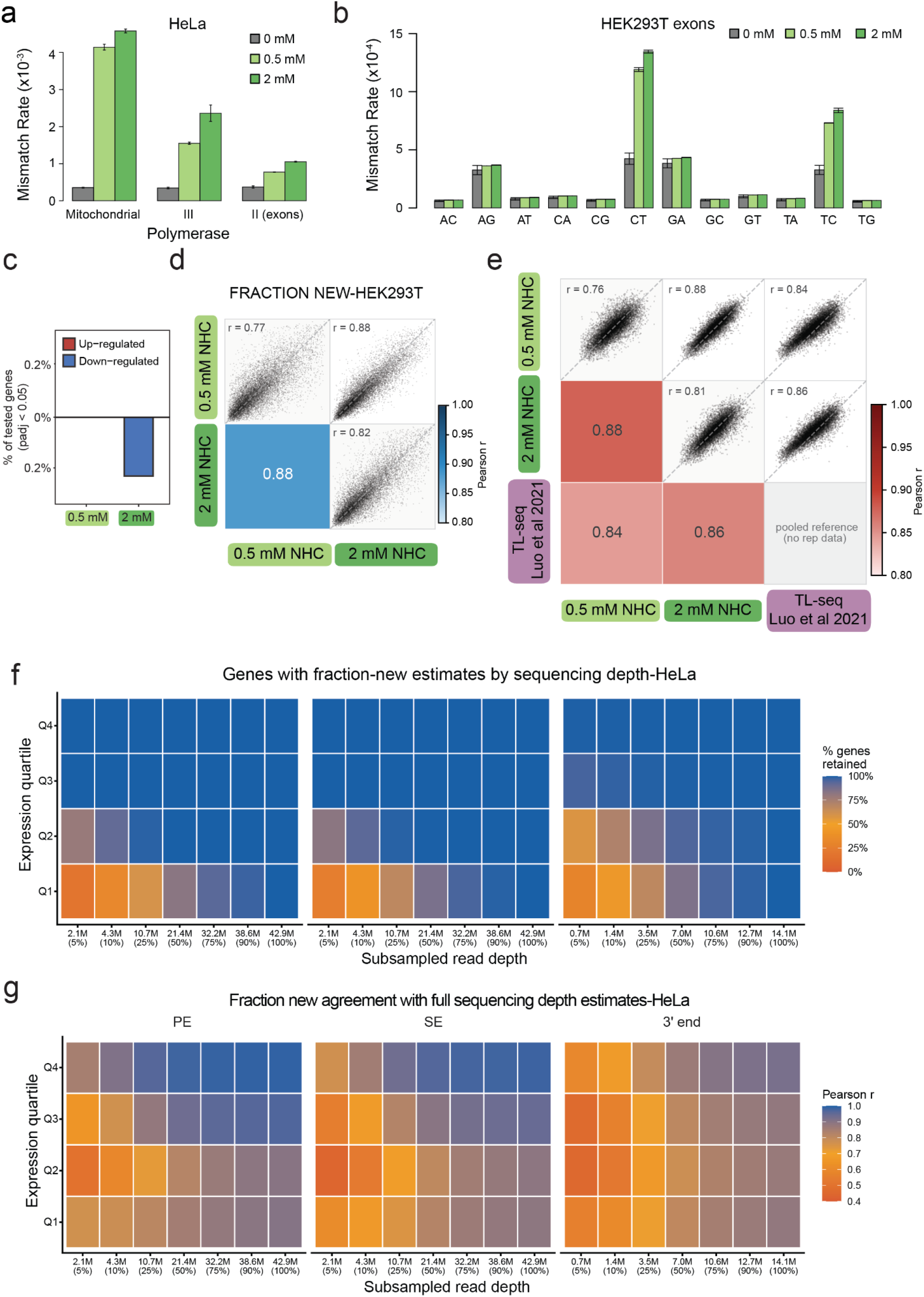
Characterizing NHC-AIR-seq performance. **a)** Mismatch rate (T-to-C and C-to-T) for HeLa cells by polymerase (mean ± range, n = 2 replicates). **b)** As in **Fig 1c**, for HEK293T, mismatch rates for all 12 possible single-nucleotide substitution types in exonic reads (mean ± range, n = 2 replicates). **c)** Differential gene expression in HEK293T at 0.5 and 2 mM NHC vs. unlabeled control (2 h), DESeq2 (Wald test; log2 fold-change shrinkage, ‘normal’ prior; padj < 0.05, Benjamini-Hochberg FDR correction), shown as percentage of tested genes up- and down-regulated per dose. 0.5 mM: n=0 significant of 16,715 tested (values below 0.1% not visible). 2 mM: n=0 up, n=20 down (100% of significant genes down-regulated) of 8,613 tested. **d)** Pairwise agreement in fraction new between NHC doses (0.5 and 2 mM) in HEK293T cells. Lower triangle, Pearson r (replicates pooled); diagonal, replicate A vs B, n=10,919 genes (shared read depth > 100 in all samples). **e)** As in (d), for RNA half-life, with Luo et al.^10^ (TimeLapse-seq, HEK293T, 2 hr 500 µM 4sU) for comparison (n=10,210 genes with shared read depth > 100 in all samples and half-lives between 0.1 and 100 hr). **f)** Fraction of genes with a valid fraction-new estimate relative to full sequencing depth, across downsampled read depths and expression quartiles (quartiles defined from full depth paired-end data; Methods), for paired-end, single-end and 3′end libraries in HeLa cells. g) As in (f), Pearson correlation of downsampled fraction-new estimates with the full-depth reference (Fisher z-averaged across 5 downsampling replicates per depth, with degrees-of-freedom weighted; Methods).

**Supplementary Figure 2:**
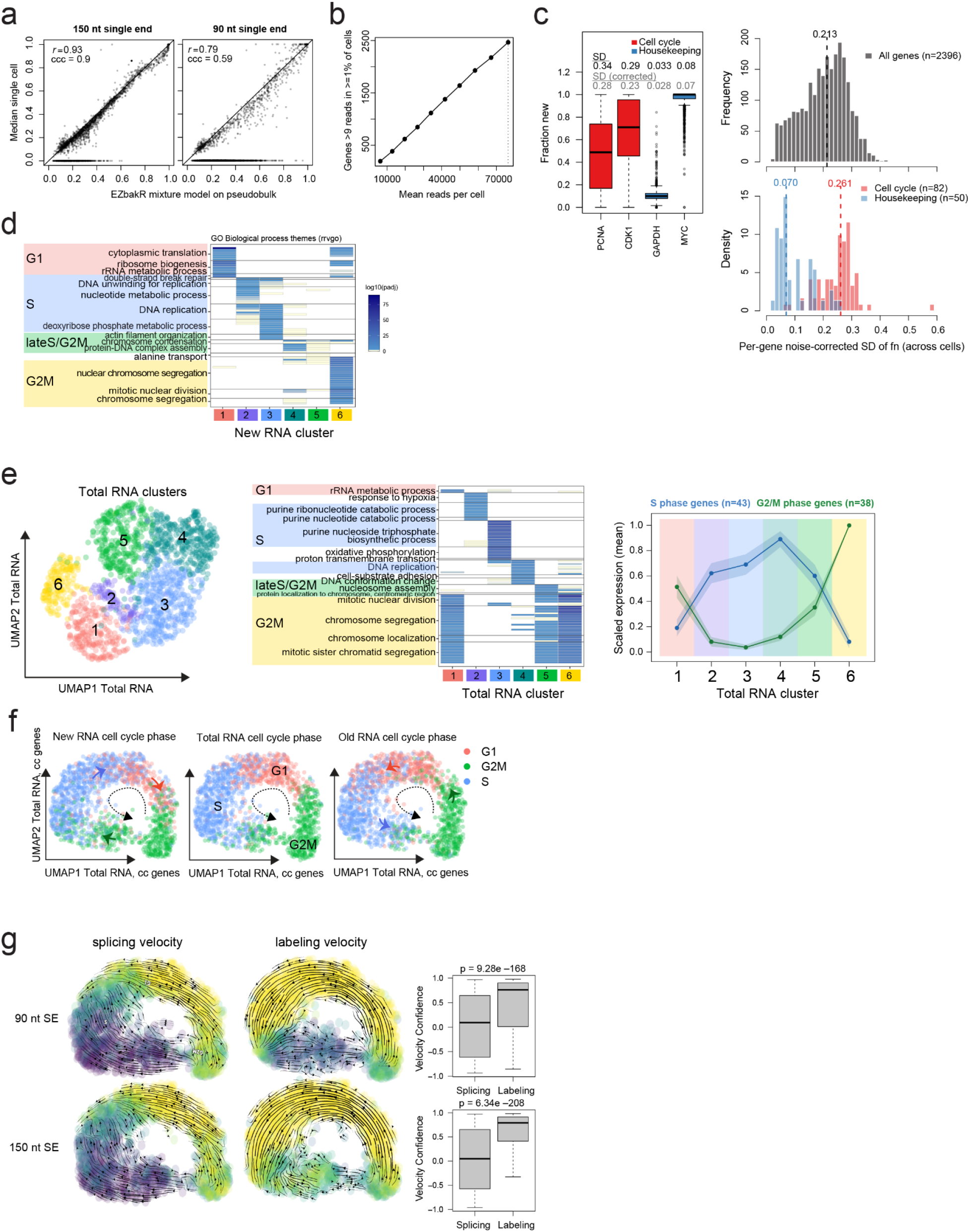
Single-cell NHC-AIR-seq reports on cell-cycle RNA dynamics. **a)** Median single-cell fraction-new estimates after trimming reads to 150 nt or 90 nt compared with EZbakR fraction-new estimates from pseudobulk data. **b)** Downsampling analysis: number of genes expected to pass the single-cell expression threshold, defined as >9 reads in at least 1% of cells, as a function of mean reads per cell in NHC-treated cells (1610 cells). Dotted line shows the observed sequencing depth in the current experiment. **c)** Left, Fraction-new distributions for representative cell-cycle genes and housekeeping genes across single cells. Raw and noise-corrected standard deviation (SD) is shown. Right, distribution of per-gene fraction-new SD corrected for binomial sampling noise to account for read coverage differences between gene sets. NAs values were dropped, and zeros were retained; dashed lines mark median values. **d)** GO biological-process enrichment themes for markers of New RNA clusters, summarized with rrvgo and grouped by inferred cell-cycle phase. Blocks for individual terms within each parent term are retained, and their color indicates -log10 adjusted P value, where values < 3 are shown in a yellow gradient, and values >= 3 are shown on a blue gradient. **e)** The top 2,000 most variable genes in the New RNA assay were identified from log-normalized counts. Expression values were scaled and used as input to PCA; the first 20 principal components were used to construct a shared nearest-neighbor graph, and cells were embedded by UMAP. Middle, GO themes for markers of Total RNA clusters. Right, mean scaled expression of S-phase and G2M-phase genes in Total RNA clusters. **f)** Cell-cycle phase assignment from Seurat CellCycleScoring using New RNA, Total RNA, or Old RNA colored on the Total RNA cell-cycle UMAP. Arrows indicate the shifted onset of phase-specific programs across new, total, and old RNA along the inferred cell-cycle trajectory. **g)** RNA velocity as in Figure 2f, but applied to 150 nt or 90 nt shortened reads.

